# Ensuring scientific reproducibility in bio-macromolecular modeling via extensive, automated benchmarks

**DOI:** 10.1101/2021.04.04.438423

**Authors:** Julia Koehler Leman, Sergey Lyskov, Steven Lewis, Jared Adolf-Bryfogle, Rebecca F. Alford, Kyle Barlow, Ziv Ben-Aharon, Daniel Farrell, Jason Fell, William A. Hansen, Ameya Harmalkar, Jeliazko Jeliazkov, Georg Kuenze, Justyna D. Krys, Ajasja Ljubetič, Amanda L. Loshbaugh, Jack Maguire, Rocco Moretti, Vikram Khipple Mulligan, Phuong T. Nguyen, Shane Ó Conchúir, Shourya S. Roy Burman, Shannon T. Smith, Frank Teets, Johanna KS Tiemann, Andrew Watkins, Hope Woods, Brahm J. Yachnin, Christopher D. Bahl, Chris Bailey-Kellogg, David Baker, Rhiju Das, Frank DiMaio, Sagar D. Khare, Tanja Kortemme, Jason W. Labonte, Kresten Lindorff-Larsen, Jens Meiler, William Schief, Ora Schueler-Furman, Justin Siegel, Amelie Stein, Vladimir Yarov-Yarovoy, Brian Kuhlman, Andrew Leaver-Fay, Dominik Gront, Jeffrey J. Gray, Richard Bonneau

**Author notes:** these authors contributed equally to this work (Julia Koehler Leman, Sergey Lyskov, Steven Lewis).

## Abstract

Each year vast international resources are wasted on irreproducible research. The scientific community has been slow to adopt standard software engineering practices, despite the increases in high-dimensional data, complexities of workflows, and computational environments. Here we show how scientific software applications can be created in a reproducible manner when simple design goals for reproducibility are met. We describe the implementation of a test server framework and 40 scientific benchmarks, covering numerous applications in Rosetta bio-macromolecular modeling. High performance computing cluster integration allows these benchmarks to run continuously and automatically. Detailed protocol captures are useful for developers and users of Rosetta and other macromolecular modeling tools. The framework and design concepts presented here are valuable for developers and users of any type of scientific software and for the scientific community to create reproducible methods. Specific examples highlight the utility of this framework and the comprehensive documentation illustrates the ease of adding new tests in a matter of hours.

## Introduction

Reproducibility in science is a systemic problem. In a survey published by *Nature* in 2016, 90% of scientists responded that there is a reproducibility crisis^1^. Over 70% of the over 1,500 researchers surveyed were unable to reproduce another scientist’s experiments and over half were unable to reproduce their own experiments. Another analysis published by *PLOS One* in 2015 concluded that, in the US alone, about half of preclinical research was irreproducible, amounting to a total of about $28 billion being wasted per year^2^!

Reproducibility in biochemistry lab experiments remains challenging to address, as it depends on the quality and purity of reagents, unstable environmental conditions, and accuracy and skill with which the experiments are performed. Even small changes in input and method ultimately lead to an altered output. In contrast, computational methods should be inherently scientifically reproducible since computer chips perform computations in the same way, removing some variations that are difficult to control. However, in addition to poorly controlled computing environment variables, computational methods become increasingly complex pipelines of data handling and processing. This effect is further compounded by the explosion of input data through “big data” efforts and exacerbated by a lack of stable, maintained, tested and well-documented software, creating a huge gap between the theoretical limit for scientific reproducibility and the current reality^3^.

These circumstances are often caused by a lack of best practices in software engineering or computer science^4,5^, errors in laboratory management during project or personnel transitions, and a lack of academic incentives for software stability, maintenance, and longevity^6^. Shifts in accuracy can occur when re-writing functionality or when several authors work on different parts of the codebase simultaneously. An increase in complexity of scientific workflows with many and overlapping options and variables can prevent scientific reproducibility, as can code implementations that lack or even prevent suitable testing^4^. Absence of testing and maintenance cause software erosion (also known as *bit rot*), leading to a loss of users and often the termination of a software project. Further, barriers are created through intellectual property agreements, competition, and refusal to share inputs, methods and detailed protocols.

As an example, in 2011 the Open Science Collaboration in Psychology tried to replicate results of 100 studies as part of the Reproducibility Project^7^. The collaboration consisting of 270 scientists could only reproduce 39% of study outcomes. Since then, some funding agencies and publishers have implemented data management plans or standards to improve reproducibility^8–11^, for instance the FAIR data management principles^12^. Guidelines to enhance reproducibility^13^ are outlined in Table 3 and are discussed in detail in an excellent editorial^14^ describing the *Ten Year Reproducibility Challenge*^15^ that is published in its own reproducibility journal ReScience C^16^. Other efforts focus directly on improving the methods with which the researchers process their data – for instance the Galaxy platform fosters accessibility, transparency, reproducibility, and collaboration in biomedical data analysis and sharing^13^.

Reproducibility is also impacted by *how* methods are developed. Comparing a newly developed method to established ones, or an improved method to a previous version, is important to assess its accuracy and performance, monitor changes and improvements over time and evaluate the cost/benefit ratio for software products to commercial entities. However, biases in publishing positive results or improvements to known methods, in conjunction with errors in methodology or statistical analyses^17^, lead to an acute need to test methods via third parties. Often, methods are developed and tested on a specific benchmark set created for that purpose and will perform better on that dataset than methods not trained on that particular dataset. A rigorous comparison and assessment require the benchmark to be independently created from the method, which unfortunately is rarely the case. Compounding issues are lack of diversity in the benchmark set (towards easier prediction targets) and reported improvements smaller than the statistical variation of the predicted results. Guidelines on how to create a high-quality benchmark^18,19^ are outlined in Table 3.

Scientific reproducibility further requires a stable, maintainable and well-tested codebase. Software testing is typically achieved on multiple levels^4,20^. Unit tests check for scientific correctness of small, individual code blocks, integration tests check an entire application by integrating various code blocks, and profile and performance tests ensure consistency in runtime and program simplicity. Scientific tests or benchmarks safeguard the scientific validity and accuracies of the predictions. They are typically only carried out during or after the development of a new method (*‘static benchmarking’*), as they require domain expertise and rely on vast computational resources to test an application on a larger dataset. However, accuracy and performance of a method depend on the test set, the details of the protocol (i.e. specific command lines, options and variables), and the software version. To overcome the static benchmarking approach, blind prediction challenges such as the Critical Assessments in protein Structure Prediction (CASP^21^), PRediction of protein Interactions (CAPRI^22^), Functional Annotation (CAFA^23^), Genome Interpretation (CAGI^24^), RNA Puzzles^25^, and Continuous Automated Model EvaluatiOn (CAMEO^26 15^) hold double-blind competitions at regular intervals. While these efforts are valuable to drive progress in method development in the scientific community, participation often requires months of commitment and does not address the reproducibility of established methods over time.

The Rosetta macromolecular modeling suite^27,28^ has been developed for over 20 years by a global community with now hundreds of developers at over 70 institutions^4,29^. This history and growth required us to adopt many best practices in software engineering^4,28^, including the implementation of a battery of tests. A detailed description of our community, including standards and practices, is available in^4^. Scientific tests are important to maintain prediction accuracies for our own community and our users (including commercial users whose licensing fees, in our case, support much of Rosetta’s infrastructure and maintenance). We further want to directly and quickly compare different protocols and implementations and monitor the effect of score function changes onto the prediction results. For many years, Rosetta applications^30,31^ and score functions^32–35^ have been tested independently using the static benchmarking approach^19,36^, often with complete protocol captures^37,38^. The disadvantage of static benchmarking is that the results become outdated due to the lack of automation. Reproducibility becomes impossible due to lack of preservation of inputs, options, environment variables and data analyses over time.

This background highlights the challenges in rigorously and continuously testing how codebase changes affect the scientific validity of a prediction method, while maintaining or improving scientific reproducibility. Running scientific benchmarks continuously (1) suffers from a lack of incentive to set up as the maintenance character of these tests collides with academic goals; (2) requires both scientific and programming/technical expertise to implement, interpret and maintain; (3) is difficult to interpret with pass/fail criteria; and (4) requires a continuous investment of considerable computational resources. Here we address these challenges by introducing a general framework for continuous scientific benchmarks for a large and increasing number of protocols in the Rosetta macromolecular modeling suite. We present the general setup of this framework, demonstrate how we solve each of the above challenges and present the results of the individual benchmarks in the supplement of this paper, complete with detailed protocol captures. The results can be used as a baseline by anyone developing macromolecular modeling methods, and the code of this framework is sufficiently general to be integrated into other types of software.

## Results and Discussion

Over the past 15 years, the Rosetta community has created its own custom-built test server framework connected to a dedicated High-Performance Computing (HPC) cluster - its setup is shown in FIG 1A and also described in the Supplement. The scientific testing setup is integrated into this framework.

**Fig 1:**
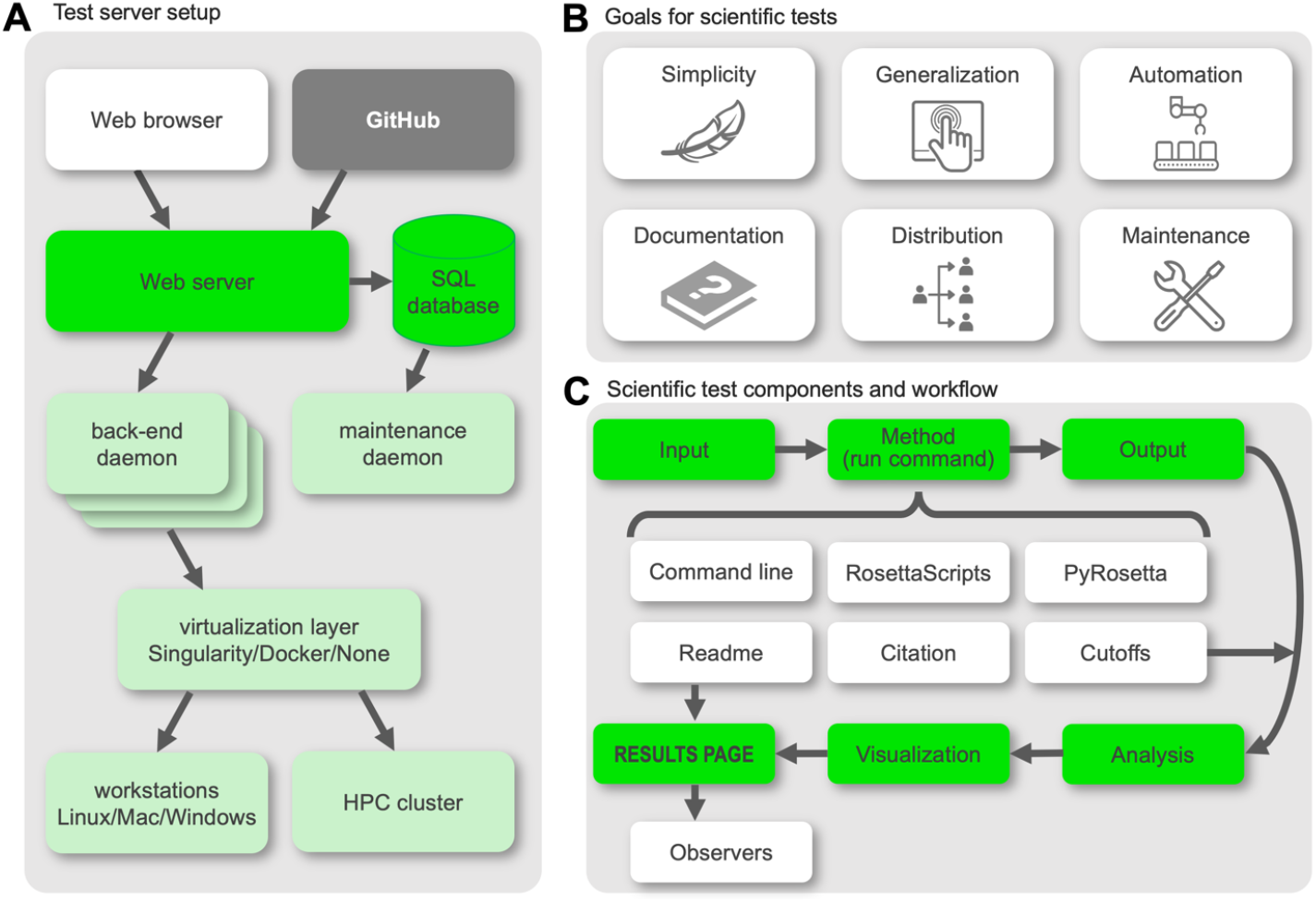
(A) Test server setup with the web browser as the user interface, the frontend in bright green and the backend in light green. The code is stored in GitHub, shown in dark gray. (B) Specific goals for our scientific tests, driven by flaws in a previous iteration of these tests. Each point is described in detail in the text. (C) Basic infrastructure of the scientific test framework, motivated by simplicity. Each box represents a file, folder, or script that is either provided in the template folder or generated throughout the protocol run. The basic workflow is highlighted in green with components that facilitate documentation and maintenance shown in white.

### Insights from previous round of scientific tests led to specific goals

The Rosetta community learned valuable lessons from the long-term maintenance (or lack thereof) of several scientific benchmark tests set up over 10 years ago. Their deterioration and development life cycle motivated specific goals that we think lead to more durable scientific benchmarks (FIG 1B): (1) Simplicity of the framework to encourage maintenance and support; (2) Generalization to support all user interfaces to the Rosetta codebase (command line, RosettaScripts^39^, PyRosetta^40,41^); (3) Automation to continuously run the tests on an HPC cluster with little manual intervention; (4) Documentation on how to add tests and scientific details of each test to allow maintenance by anyone with a general science or Rosetta background; (5) Distribution of the tests to both the Rosetta community and their users, and publicizing their existence to encourage addition of new tests and maintenance by the community; (6) Maintenance of the tests, facilitated by each of the previous points.

### Simplicity: Simple setup facilitates broad adoption and support from our community

To encourage our community to contribute as many tests as possible, the testing framework needs to be simple and support fast and easy addition of tests. We decided on a Python framework that integrates well with our pre-existing testing HPC cluster. We further require these tests to be able to run on local machines (with different operating systems) as well as various HPC clusters with minimal adjustments. Debugging the scripts should be as simple as possible. With these requirements in mind, we decided on a setup as shown in FIG 1C. We provide a template directory with all necessary files (described in detail in *Methods*). Simple modifications like naming scripts in the order in which they run - e.g. *0.compile.py* to *9.finalize.py* - greatly facilitates debugging or extension by new users.

### Generalization: New tests support interfaces of command line, PyRosetta, or RosettaScripts

Rosetta supports several interfaces to facilitate quick protocol development while lowering the necessary expertise required by new developers to join our community^4^. Many mainstream protocols have been developed as standalone applications to be run via command line, while customized protocols have been developed in RosettaScripts^39^ and PyRosetta^40,41^. For our test server framework, we sought a general code design that allows input from all three interfaces while supporting different types of outputs, quality measures, and analyses, sometimes even written in different scripting languages.

### Automation: Tests require substantial compute power and are run on a dedicated test server

Running scientific benchmarks requires extensive CPU time; hence we chose to integrate them with our own custom-built test server framework connected to a dedicated HPC cluster (FIG 1A and Supplement). This test server framework consists of two main components: the backend holds low-level primitive code for compilation on different operating systems and HPC environments, cluster submission scripts and web server integration code. The frontend contains the test directories that are implemented by the test author. Our test server is accessible through a convenient web interface (FIG. 2A; available at https://benchmark.graylab.jhu.edu/). This framework has had a hugely positive impact on the growth and maintenance of both the Rosetta software and our community, due to its accessibility, GitHub integration, ease-of-use, and automation. In software communities that lack the ability to set up a dedicated test server, integration testing via external services like Travis CI or Jenkins are an alternative.

**Fig 2:**
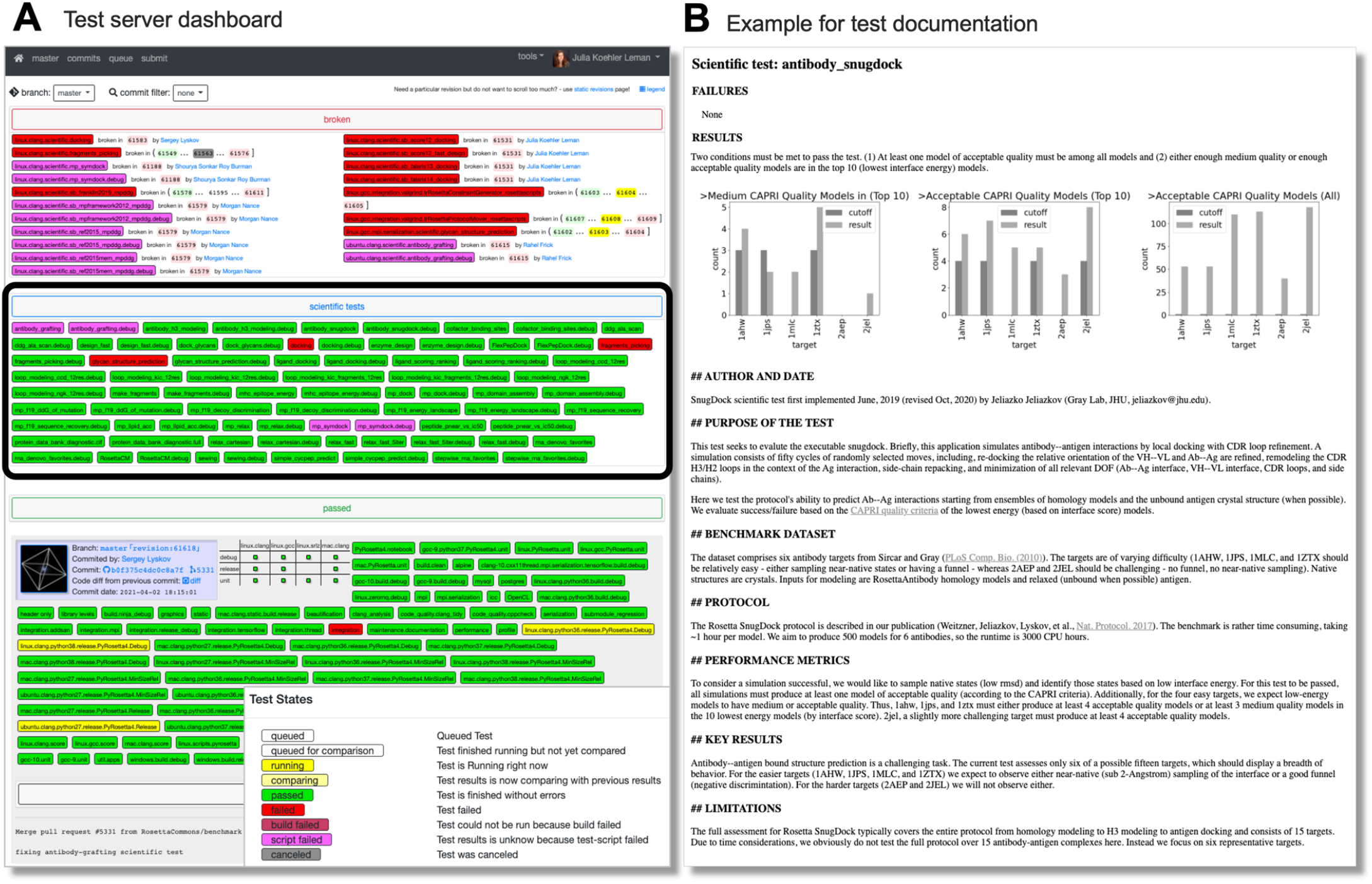
(A) Dashboard of our benchmark server testing infrastructure. Each test is colored according to its test results: red denotes breakage, magenta denotes script failure, green denotes passing of a test, yellow denotes the test is currently running, and white denotes the test has yet to be run. All broken tests are shown prominently at the top of the page. All scientific tests are shown in the blue tab below (also encircled in bold black). Tests of the latest revision merged into the main branch are shown below with information about the committer, the pull request ID, a link to the code difference, and the commit message. (B) The results page that shows the results of the run, the documentation, and the description of whether the test passes or fails. Results pages are automatically generated at the end of the run for each test.

The RosettaCommons supports our benchmarking effort through expansion of our centralized test server cluster hardware and labor with an annual budget (see Supplement and reference ^4^). Because the scientific tests are integrated into our test server framework, authors of the tests can focus on the scientific protocols (starting from a template directory set up as in FIG 1C) instead of debugging errors in compilation, cluster submission and computational environment. This pattern also makes these tests system-independent (the author writes the setup for a local machine and runs it on this server), i.e., portable between operating systems and computational environments. We currently limit the runtime per scientific test to typically 1,000-2,000 CPU hours.

Due to the required computational resources, we are unable to test every code revision in the main development branch of Rosetta; instead, we dedicate computational nodes to the scientific tests and run tests such that the nodes are continuously occupied. We found that scheduling the earliest-run test on an individual rolling basis, as compute nodes become available, is most efficient in balancing the server load while keeping nodes available for tests in feature branches. Upon discovery of a test failure and to find the specific revision (and therefore the code change) that caused the failure, our *bisect* tool schedules intermediate revisions on a low-priority basis. All test results are stored in the database and are accessible through a web interface (FIG 2).

### Documentation: Anyone can quickly and easily add new tests

Creating well designed scientific benchmarks requires expertise in defining the scientific objective, establishing a protocol, and creating a high-quality test dataset. The last step of incorporating the test into our framework should be as simple as possible (as per our *simplicity* requirement). Once the dataset, interface (command line, RosettaScripts, or PyRosetta), specific command line, and quality measures have been chosen, the author can simply follow the individual steps outlined on the documentation page^42^ to contribute the test; the template guides the setup. We found that the setup is simple enough that untrained individuals can contribute a test in a few hours based on documentation alone – hence we achieved our goal of simplicity and detail in our documentation.

One of the reasons for deterioration of earlier scientific tests was lack of maintenance due to insufficient documentation. Our goal is to drive the creation of extensive documentation for each test such that anybody with an average scientific knowledge of biophysics and introductory knowledge of programming in Rosetta can understand and maintain the tests. To ensure comprehensive documentation and consistency between tests, we provide a readme template with specific sections and questions that need to be answered for each test (see Supplement). The template discourages writing short, insufficient, free-form documentation, and instead encourages the addition of important details and significantly lowers the barrier for writing extensive documentation. The questionnaire-style readme template saves time to locate necessary details to repair broken tests. The extent and quality of documentation is independently approved by a pull-request reviewer before the test is merged into the main repository. The benchmark framework is configured such that documentation needs to be written once and is then directly embedded into the results page. Thus, the documentation is accessible both in the code and on the web interface while eliminating text duplication that could lead to discrepancies and confusion.

### Distribution: Additions and usage of tests by our community requires broad distribution

Earlier scientific tests also deteriorated due to poor communication as to the existence of these tests, which resulted in a small pool of maintainers. Because our new scientific tests are integrated into our test server framework which the majority of our community uses and monitors, developers are immediately aware of the tests that exist and their pass/fail status. In conjunction with regular announcements to our community, this visibility should significantly broaden the number of people able and willing to sustain the scientific tests for a long time. If we nevertheless find that our new tests deteriorate, we will host a hackathon (eXtreme Rosetta Workshop^4^) to supplement or repair these tests in a concentrated effort.

### Maintenance: Test failures are handled by a defined procedure

The often overlooked, *real* work in software development is not necessarily the development of the software itself, but its maintenance. We have a system in place outlining how test failures are handled and by whom. Each test has at least one dedicated maintainer (aka ‘observer’, usually the test author) who is notified of the test breakage via email and whose responsibility it is to repair the test. Test failures can be three-fold: technical failures, stochastic failures, or scientific failures. Technical failures (such as compiler errors, script failures due to new versions of programs, etc.) typically require small adjustments and fall under the responsibility of the test author and our dedicated test engineer.

Stochastic failures are an uncommon feature in software testing but are possible in this framework. Rosetta often uses Metropolis Monte Carlo algorithms and thus has an element of randomness present in most protocols. Scientific tests are interpreted in a Boolean pass/fail fashion but generally have an underlying statistical interpretation and are sampling from a distribution against a chosen target value. The randomness of Monte Carlo will occasionally cause a stochastic test failure because those runs happen to produce poor predictions by the tested metric. This is handled by simply rerunning the test: rare stochastic failures are not likely to occur repeatedly, and if they do, it merits re-examination of the test to change its structure or pass/fail criteria.

A scientific failure requires more in-depth troubleshooting and falls under the responsibility of the maintainer. If the maintainer does not fix the test, we have a rank-order of responsibilities to enforce the test repair. The principal investigator of the test designates someone in their lab. If the necessary expertise does not exist in the lab at the time (usually because people have moved on in their career), repairing the test becomes the responsibility of the person who broke it. If this developer lacks the expertise, the repair becomes community responsibility, which typically falls onto one of our senior developers.

### Most major Rosetta protocols are now implemented as scientific benchmarks

Using the framework described above, our community implemented 40 scientific benchmarks spanning a broad range of applications including antibody modeling, docking, loop modeling, incorporation of NMR data, ligand docking, protein design, flexible peptide docking, membrane protein modeling etc. (Table 1 and Supplement). Each benchmark is unique in its selection of targets in the benchmark set, the specific protocol that is run, the quality metrics that are evaluated, and the analysis to define the pass/fail criterion. The details for all of the benchmarks are provided in the comprehensive supplement to this paper. We further publish the benchmarks with results and protocol captures on our website (https://graylab.jhu.edu/download/rosetta-scientific-tests/) twice per year for our users to see, download, run, and compare their own methods against. This transparency is crucial for representation of realistic performance and to enhance scientific reproducibility of our tools.

**Table 1:** Scientific tests continuously running on our testing server framework. The number of tests is constantly being expanded. The test suite is the overall application, the test is the specific test, implemented by the test author(s). The quality measures are evaluated to choose a pass/fail criterion. The targets are the number of different proteins (or biomolecules) tested on, nstruct is the number of models built for each target, and the runtime in CPU hours is the total runtime over all targets.

| test suite | tests | ref | test author | quality measures | targets | nstruct | runtime in CPUh |
| --- | --- | --- | --- | --- | --- | --- | --- |
| antibodies | antibody_grafting | 43 | Jeliazko Jeliazkov | fraction residues within rmsd to native | 48 | 1 | 3 |
|  | antibody_h3_modeling | 44 |  | score vs rmsd | 6 | 500 | 3000 |
|  | antibody_snugdock | 45 |  | l_sc vs l_rmsd | 6 | 500 | 3000 |
| carbohydrates | dock_glycans | 46 | Jason Labonte | l_sc vs l_rmsd | 4 | 500 | 60 |
|  | glycan_structure_prediction | 47 | Jared Adolf-Bryfogle | score vs rmsd | 4 | 500 | 950 |
| comparative modeling | RosettaCM | 48 | Jason Fell | GDT-MM | 16 | 200 | 1800 |
| design | ddg_alanine_scan | 49 | Ajasja Ljubetič | R, MAE, fraction correctly classified | 19: 381 | 1 | 3 |
| design | SEWING | 50 | Frank Teets | MotifScorer, InterModelMotifScorer | 1 | 100 | 75 |
| design | enzyme_design | 51 | Rocco Moretti | various sequence recoveries | 50 | 1 | 50 |
| design | design_fast | 52 | Jack Maguire, Chris Bahl | score vs seqrec | 48 | 100 | 2600 |
| design, interfaces | cofactor_binding_sites | 58 | Amanda Loshbaugh | rank top, position profile similarity | 7 | 200 | 170 |
| design, immune system | mhc_epitope_energy | 53 | Brahm Yachnin | degree of de-immunization, among others | 50 | 100 | 2000 |
| docking | protein_protein_docking | 54 | Shourya SR Burman, | l_sc vs l_rmsd | 10 | 1000 | 150 |
|  | ensemble docking | 55 | Ameya Hamalkar | l_sc vs l_rmsd | 3 | 1000 | TBD |
| FlexPepDock | FlexPepDock | 56 | Ziv Ben-Aharon | reweighted l_sc vs backbone l_rmsd | 2 | 200 | 70 |
| fragments | fragment_picking | 57 | Justyna Kryś, Dominik Gront | rmsd | 10 | 400 | 2000 |
| fragments | make fragments pipeline | 57 | Daniel Farrell | coverage, precision | 65 | 1 | 3000 |
| ligand docking | ligand_docking | 59 | Shannon Smith | delta_lsc vs ligand_rmsd | 50 | 200 | 2000 |
|  | ligand_scoring_ranking | 59 |  | Spearman and Pearson correlation coefficient | 57: 285 | 1 | 2 |
| loop modeling | loop_modeling_CCD | 60 | Phuong Tran,<br>Shane Ó Conchúir | score vs loop_rmsd | 7 | 500 | 500 |
|  | loop_modeling_KIC | 61 |  | score vs loop_rmsd | 7 | 500 | 620 |
|  | loop_modeling_KIC_fragments | 62 |  | score vs loop_rmsd | 7 | 500 | 760 |
|  | loop_modeling_NGK | 63 |  | score vs loop_rmsd | 7 | 500 | 570 |
| membrane protein energy function | mp_f19_energy_landscape | 36 | Rebecca Alford | ddG, depth and title angle | 4 | 1 | 10 |
|  | mp_f19_decoy_discrimination | 36 |  | score vs rmsd, Wrms | 4x100 | 1 | 2000 |
|  | mp_f19_sequence_recovery | 36 |  | sequence recovery, Kullback-Leibler divergence | 130 | 1 | 500 |
|  | mp_f19_ddG_of_mutation | 64 |  | Pearson correlation coefficient | 3 | 1 | 1 |
| membrane proteins | mp_dock | 65 | Julia Koehler Leman,<br>Rebecca Alford | l_sc vs l_rmsd | 10 | 1000 | 200 |
|  | mp_domain_assembly | 66 |  | score vs rmsd | 5 | 5000 | 700 |
|  | mp_lipid_acc | 67 |  | accuracy | 223 | 1 | 2 |
|  | mp_relax | 65 |  | score vs rmsd | 4 | 100 | 40 |
|  | mp_symdock | 65 |  | l_sc vs rmsd | 5 | 1000 | 140 |
| PDB diagnostic | PDB_diagnostic | NA | Steven Lewis, William Hansen, Sergey Lyskov | read-in error type | entire PDB | 1 | 1000 |
| peptide structure prediction | simple_cycpep_predict | 68 | Vikram K. Mulligan | score vs rmsd, PNear | 1 | ~800,000 | 320 |
|  | peptide_pnear_vs_ic50 | 69 |  | IC50 vs folding energy | 7 | 80,000 | 400 |
| refinement | relax_cartesian | 31 | Julia Koehler Leman | score vs rmsd | 12 | 100 | 120 |
|  | relax_fast | 70 |  | score vs rmsd | 12 | 100 | 120 |
|  | relax_fast_5iter | 70 |  | score vs rmsd | 12 | 100 | 120 |
| RNA | rna_denovo_favorites | 71 | Andy Watkins | score vs rmsd | 12 | 200 | 120 |
|  | stepwise_rna_favorites | 72 |  | score vs rmsd | 12 | 200 | 240 |

for table references – not part of the actual manuscript:
antibody grafting <sup>43</sup>, antibody H3 <sup>44</sup>, antibody snugdock <sup>45</sup>, dock glycans <sup>46</sup>, glycan structure prediction <sup>47</sup>, RosettaCM <sup>48</sup>, ddg\_alanine\_scan <sup>49</sup>, SEWING <sup>50</sup>, enzyme design <sup>51</sup>, fast design <sup>52</sup>, mhc epitopes <sup>53</sup>, docking <sup>54</sup>, ensemble docking <sup>55</sup>, FlexPepDock <sup>56</sup>, fragment picking <sup>57</sup>, make fragments <sup>57</sup>, [iHyb], cofactors <sup>58</sup>, ligand docking <sup>59</sup>, ligand scoring ranking <sup>59</sup>, loops CCD <sup>60</sup>, loops KIC <sup>61</sup>, loops KIC fr <sup>62</sup>, loops NGK <sup>63</sup>, mp energies <sup>36</sup>, mp decoy discrimination <sup>36</sup>, mp seq rec <sup>36</sup>, mp ddg <sup>64</sup>, mp dock <sup>65</sup>, mp domain assembly <sup>66</sup>, mp lipid acc <sup>67</sup>, mp relax <sup>65</sup>, mp symdock <sup>65</sup>, [abinitio (Simons, K. T., Kooperberg, 1997)], PDB diagnostics, simple cycpep <sup>68</sup>, pnear <sup>69</sup>, relax cart <sup>31</sup>, relax fast <sup>70</sup>, relax fast5 <sup>70</sup>, RNA denovo <sup>71</sup>, stepwise <sup>72</sup>

### Standardizing workflows highlights heterogeneity in score function implementations

Standardizing the workflows and creating this framework provides us with the possibility of running some protocols with different score functions. Rosetta has been developed over the past 25 years and the score function has been constantly improved over this timeframe. Details of this evolution and the latest standard score function REF2015 can be found in references ^34,35^. The attempt to easily switch score functions for an application reveals a major challenge: many applications employ the global default score function differently, a problem exacerbated by the various user interfaces to the code (see Supplement for details). The heterogeneity in implementations makes it impossible to easily test different score functions for all of the applications and reveals that it hinders both progress and unification of the score functions, possibly into a single one.

### Use case #1: Test framework allows comparison of score functions for multiple protocols

Using our framework, we are able to compare different score functions for various applications: protein-protein docking, high-resolution refinement, loop modeling, design, ligand docking, and membrane protein ddG’s (Table 2 and FIGs 3 - 5). We test the latest four score functions: score12, talaris2013, talaris2014, and REF2015 for all but ligand docking and membrane protein ddG’s. Ligand docking has a special score function and requires adjustments – we test the ligand score function, talaris2014, REF2015, and the experimental score function betaNov2016. Membrane protein ddG’s are tested on the membrane score functions mpframework2012, REF2015_mem, franklin2019 and the non-membrane score function REF2015 as a control.

**Table 2:**
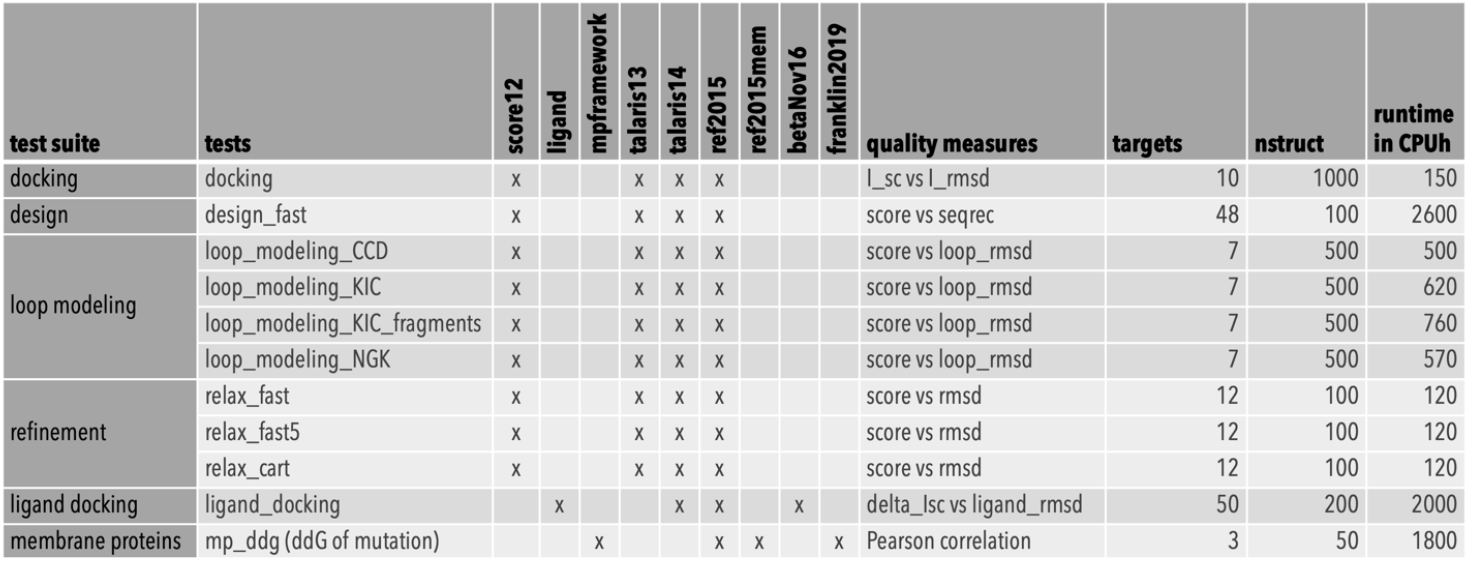
Tests for which we compare different scorefunctions (score12, talaris13, talaris14, ref2015, ligand, betaNov16, mpframework, ref2015mem, and franklin2019), complete with quality measures, number of targets in each benchmark, number of models created (nstruct) and runtime in CPU hours per scorefunction. The ligand docking and membrane ddG applications require specialized scorefunctions.

**Fig 3:**
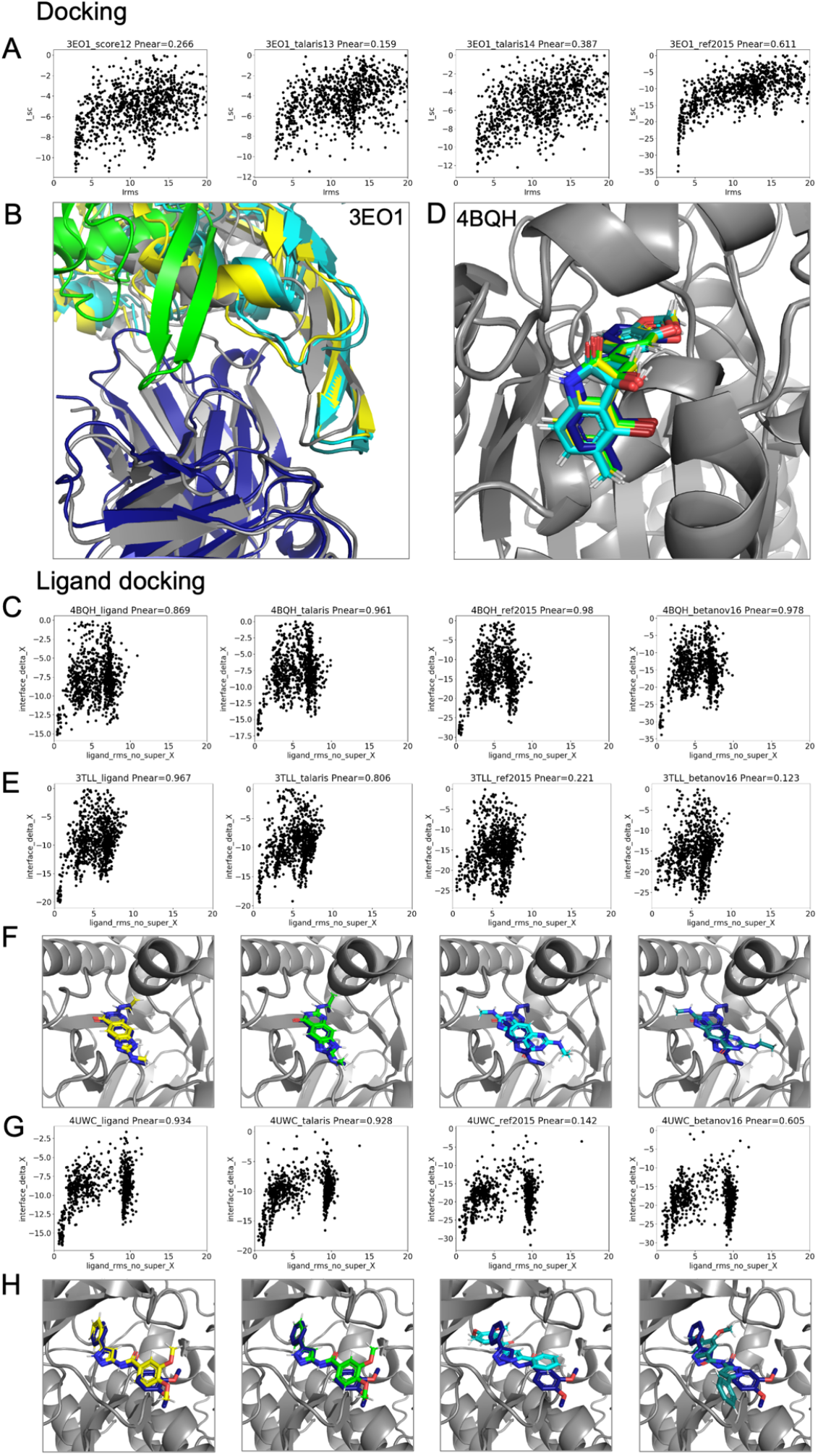
Comparison of different score functions for different applications using the P_Near_ metric as indication of “funnel quality”. P_Near_ falls between 0 (no funnel or incorrect global minimum) and 1 (perfect funnel). The lambda parameter indicates the spread on the x-axis and is set to 4.0. Score functions are sorted from oldest to newest (left to right) and the models are colored in gray as the native (PDB) structure, then according to the score functions in order: yellow, green, cyan, and teal. (A) and (B) comparison for protein-protein docking on target 3eo1. The starting model is shown in dark blue - the docking partner of the starting model is too far away to be shown in the picture. The quality of the prediction improves over different score functions as indicated by tightening of the energy funnel. (C) and (D) comparison for ligand docking on target 4bqh. The native ligand pose is shown in dark blue. The quality of the prediction improves over different score functions as indicated by tightening of the energy funnel. (E / F) and (G / H) ligand docking comparison on targets 3tll and 4uwc, respectively. Newer score functions lower the energy of an incorrect, alternative docking conformation, leading to a worse prediction.

**Fig 4:**
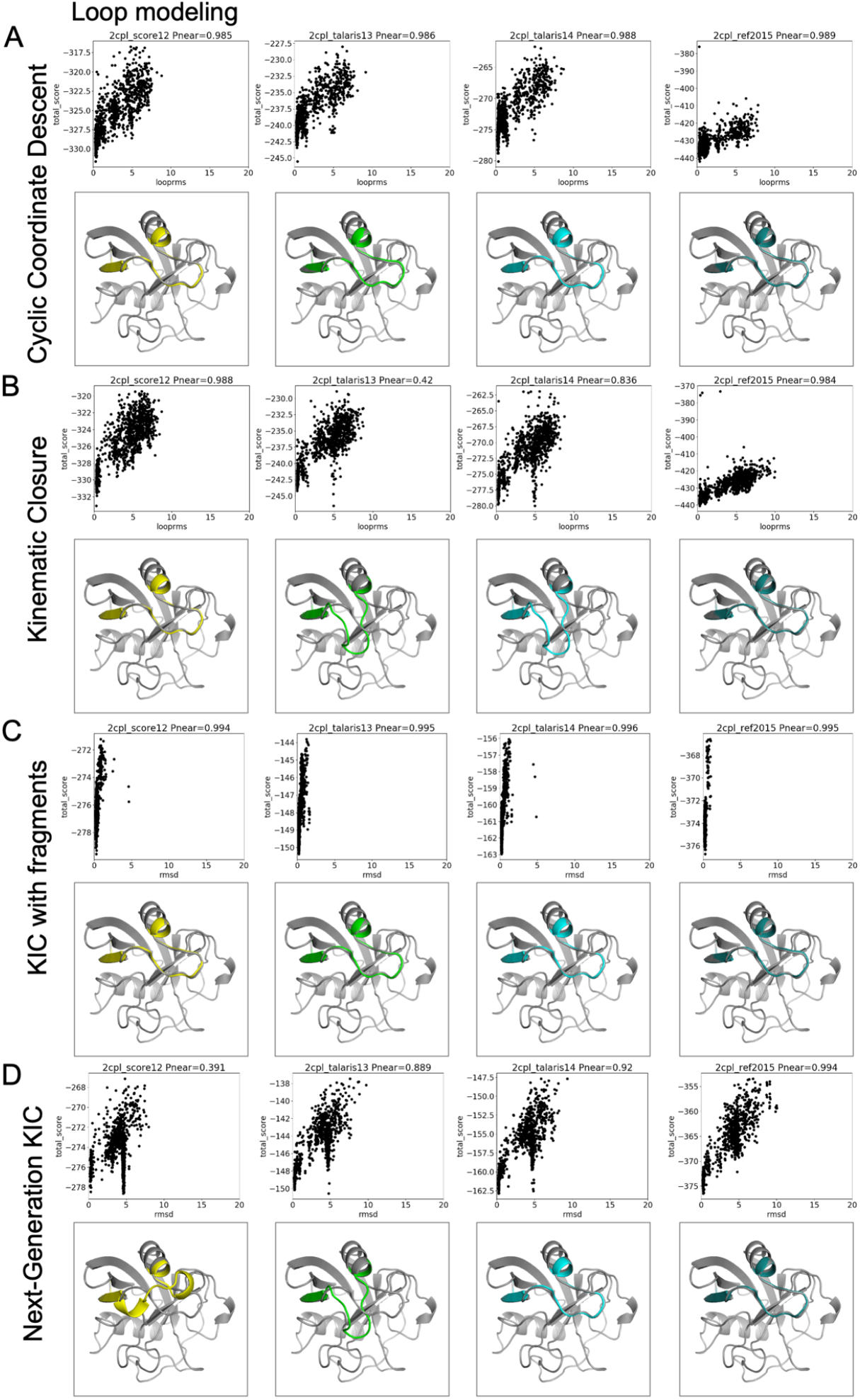
Comparison of different score functions for different loop modeling protocols. Score functions are sorted from oldest to newest (left to right) and the models are colored in gray as the native (PDB) structure, then according to the score functions in order: yellow, green, cyan, and teal. This figure shows a particular interesting example, which is not necessarily representative for other targets. Interesting for this target are the differences in the energy landscapes that are sampled and the presence of a second, incorrect conformation with low energy for some protocols and some score functions, but not others. For 3 out of 7 targets in our comparison, including this one, most conformations that KIC with fragments samples, are close to the native structure. Again, for larger benchmarks, this is likely not as often the case. The protocols are (1) Cyclic Coordinate Descent - CCD, (2) Kinematic Closure - KIC, (3) KIC with fragments, and (4) Nextgeneration KIC - NGK.

**Fig 5:**
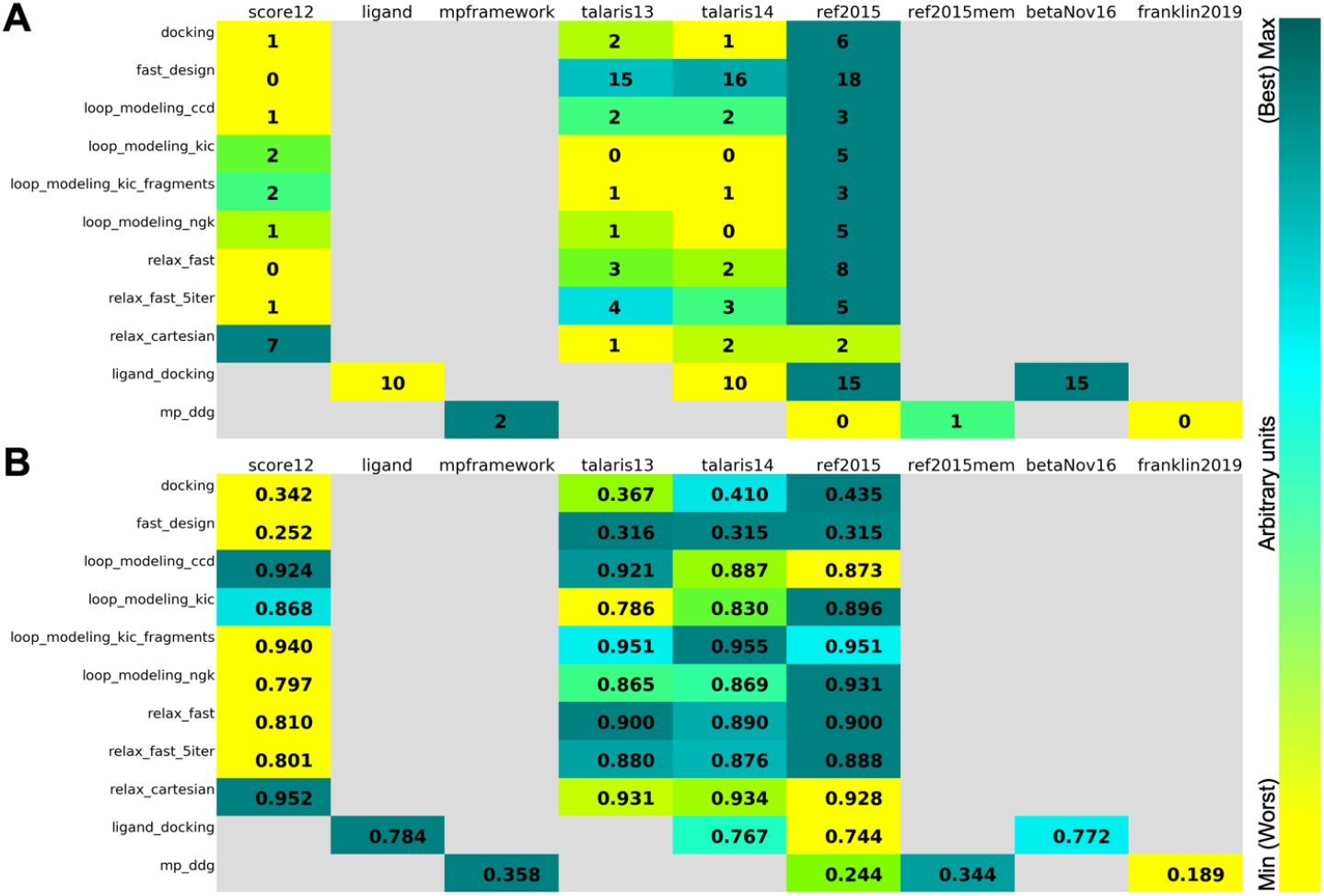
Comparison of different scorefunctions (one per column) for different applications and protocols, using the P_Near_ metric as indication of “funnel quality”. P_Near_ falls between 0 (no funnel or incorrect global minimum) and 1 (perfect funnel). The lambda parameter indicates the spread on the x-axis and is set to 4.0 in our comparison. Cells are colored according to the color bar on the right, yellow is better. Zero values in dark blue indicate unavailable data. (A) The panel uses a “winner-takes-all” comparison: for each protein, the score function with the highest (i.e., best) P_Near_ value (see methods) gets a point. Points are then summed by column, identifying the score function with the most and highest P_Near_ values across proteins, the higher the better. (B) The averages of the P_Near_ values for each score function were used, i.e. computed over each column. Higher values are better.

The benchmark sets and quality metrics are described in Table 2 and in detail in the Supplement. To compare the score functions, we plot each application’s quality metrics (for instance interface score vs. interface RMSD for protein-protein docking, total score vs. loop RMSD for loop modeling). We then evaluate the “funnel quality” by computing the P_Near_ metric, which falls between 0 and 1, with higher values indicating higher quality^68,73^. For the protein design test, we compute the average sequence similarity of the 10 lowest-scoring (best) models instead of P_Near_ and for the membrane ddG test we use the Pearson correlation coefficient between experimental and predicted ddG’s. We further summarize the quality metrics per protocol and score function by a ‘winner-takes-it-all’ comparison (FIG 5 A) and by an average metric over all targets per application per score function (FIG 5 B).

A few main observations follow from this comparison: at first glance, in this comparison, REF2015 performs generally better overall, yet the best score function to use depends on the application – even different types of protocols can impact prediction accuracy. However, it should be noted that some tests have a small sample size due to the required computational resources, therefore impacting the statistical significance of these outcomes. Second, more recent score functions are not automatically better for any given application, likely because performance depends on how the score function was developed and tested. For a more detailed discussion, see the Supplement. Third, results differ in some cases depending on how the data were summarized; the top-performing score functions per application from the ‘winner-takes-it-all’ comparison are not necessarily the top performers when the average of the P_Near_ value is used, as can be seen in ligand docking (FIG 5B – reference ^59^ discussed this in depth).

### Use case #2: Scientific test framework facilitates bug fixes and maintenance

The scientific test framework is also useful for code maintenance, to ensure that the correction of bugs does not invalidate the scientific performance of Rosetta. For example, in October 2019, we identified an integer division error in one of our core libraries: the fraction 2/3 was incorrectly assumed to evaluate to 0.6666…, when in fact integer division discards remainders, yielding 0. This calculation affected the computation of hydrogen bonding energies and their derivatives, and correcting it resulted in a small but perceptible change in some of the hydrogen bond energies. This led to the need to balance between fixing the bug and managing the complex interdependencies or to preserve the existing scoring behavior, since the rest of the score function had been calibrated with the bug present. By running the scientific tests on a development branch in which we had fixed the bug, we confirmed that although the correction results in a small change in the energies, it had no perceptible effect on the scientific accuracy of large-scale sampling runs for structure prediction, docking, design, and any other protocol tested. This allowed us to make the correction without harming Rosetta’s scientific performance. We are certain that the scientific tests will be invaluable for ensuring that future bug-fixing and refactoring efforts do not hinder the scientific performance of our software, thus illustrating a key example of scientific benchmarks informing substantive decisions developers have to make as they navigate code life cycles.

### Use case #3: Test framework allows detailed investigation of new score functions under development

Although small molecules and proteins are generally more rigid structures, intermediate-scale molecules are frequently disordered and flexible. A recent study shows that Rosetta’s estimates of rigidity (using the funnel quality metric P_Near_ computed to a designed binding conformation) for peptides designed to bind to and inhibit a target of therapeutic interest correlate well with IC_50_ values^69^. Since this prediction has relevance to computer-aided drug development efforts, we want to ensure that future protocol development would not impair these predictions. We created a test (called *peptide_pnear_vs_ic50*) which performs rigidity analysis on a pool of peptides that were previously characterized experimentally, and computes the correlation coefficient for the P_Near_ values from predicted models to the experimentally-measured IC_50_ values. We find that the current default score function, REF2015, produces much better predictions than the legacy Talaris2013 and Talaris2014 score functions (R^2^ = 0.53, 0.53, and 0.90 for Talaris2013, Talaris2014, and REF2015, respectively), indicating an improvement of the score function accuracy for this particular application^34^. However, this correlation is considerably worse with the score function Beta currently under development (R^2^ =0.19). This reveals problems in the candidate next-generation score function that will have to be addressed before it becomes the default. Our scientific tests embedded in the test server framework provide a means of rapidly benchmarking and addressing these problems.

### Use case #4: This framework and tests encourage scientific reproducibility on several levels

How is this framework useful beyond the specific tests mentioned here? Its usefulness for Rosetta developers and users lies in the protocol captures, the specific performance of each protocol and the knowledge that the scientific performance is monitored over time. Developers of macromolecular modeling methods outside of Rosetta can use and run the exact test protocol captures to compare Rosetta’s results to their own, newly developed methods. The code for the general framework to run large-scale, continuous, automated tests is available under the standard Rosetta license and is useful for developers of any type of software. Lastly, the framework highlights how software can be developed in a scientifically reproducible manner, lessons of which are useful and necessary for the scientific community at large. While we recognize the time and work required to implement such tests and the underlying framework, the benefits far outweigh the effort spent in trying to reproduce results that were implemented in a manner that lacks necessary aspects for reproducibility, as discussed in Table 3.

**Table 3:** Guidelines for reproducible research and for development of high-quality methods.

| General guidelines for reproducibility | Guidelines for high-quality benchmarks |
| --- | --- |
| 1. document artifacts<br>2. share input, output and exact workflow in detail under an open license in public repositories<br>3. cite the data, software and workflows<br>4. use persistent links in the publication<br>5. journals should check for reproducibility<br>6. funding agencies should fund reproducibility research | 1. define scientific question for the benchmark<br>2. define quality metrics that are practically relevant<br>3. diversify examples in benchmark set to cover easy and difficult targets<br>4. separate benchmark set from developed method<br>5. pick cutting edge methods to compare your method to<br>6. use benchmarked methods that are freely available |

## Conclusion

Here we present a test server framework for continuously running scientific benchmarks on an integrated HPC cluster and detail the manner in which this framework has had a positive and substantive effect on our large community of scientists. The framework itself is sufficiently general that it could in principle be used on many types of scientific software. We use it on Rosetta protocols that cover the three main interfaces to the codebase: the command line, RosettaScripts and PyRosetta. New benchmarks are easily added and debugged, and the workflow for setting them up is well-documented and general: new tests can be added in a matter of hours and require minimal coding experience in Rosetta. We provide detailed documentation and consistency in the presentation of results, thereby facilitating maintenance by more than just experts in the community and ensuring longevity of these tests. Automated and continuous runs of these tests allow us to recognize shifts in performance, as development is simultaneously carried out on several interdependent but otherwise unrelated fronts. Thus, we are able to build a longitudinal map of accuracy and scientific correctness in a constantly evolving codebase (for ourselves and our users), provide realistic protocol captures of how to run applications, and build tools that follow guidelines for improving reproducibility. Diversity in the choice of targets in the benchmark sets provides a realistic performance somewhat insulated from institutional and career incentives. So far, 40 benchmarks for various biomolecular systems and prediction tasks have been added to our server framework and more will be added over time. Running these benchmarks requires a substantial amount of resources, which are funded through RosettaCommons, since such benchmarks are a priority for software sustainability. Even though our setup involves integration of a custom software framework and web interface with typical HPC hardware, we expect our design choices to be of general interest and integrable with paid services such as Travis CI^74^ or Jenkins^75^. This framework demonstrates how challenges in scientific reproducibility can be approached and handled in a general manner, even in a large and diverse community.

## Methods

The RosettaCommons community of developers has emphasized software testing for over 15 years. To support our community of hundreds of developers, our user base of tens of thousands of users, and the codebase of over 3 million lines of code^4^, we implemented a custom testing architecture to fit our needs. We use this platform (a.k.a. the “Benchmark Server”) to run all of our tests including unit tests, integration tests, profile tests, style tests, score function tests, build tests and others. Using this benchmark server to implement scientific tests is therefore a natural extension of its current use. Our custom testing software runs on a dedicated HPC cluster (which also runs the ROSIE server^76^), paid for by the RosettaCommons from government and non-profit funding, and commercial licensing fees.

### The backend of the benchmark infrastructure consists of several servers

Our testing infrastructure consists of a number of machines:

1. ***Database server***. Our data center stores information about revisions, test and sub-test results as well as auxiliary data like comments to revisions or list of branches that are currently tracked via GitHub^4,77^. We are using PostgreSQL.
2. ***Web server***. The web interface for Rosetta developers connects to the database server. When a developer asks for a particular revision or test results, the web server gathers these data from the database server, generates the HTML page and sends it to the developer who looks at the page in a web browser. The web server also allows developers to queue new tests through the submit page on the web interface.
3. ***Revision daemon***. This application watches the state of various branches, queues tests and sends notifications. The daemon tracks the list of branches and periodically checks if a new revision for a particular branch was committed. When a new revision has been committed, it schedules the default test set for that branch. The daemon also watches for open pull requests on GitHub, and for each pull request, it checks for specific test labels (for instance ‘standard tests’). The revision daemon schedules any tests with that label for that pull-request.

Because scientific tests require an enormous amount of compute power, we are currently unable to test every single revision in the Rosetta main branch. Instead, we run scientific tests on a best-effort basis. The tests run continuously, but because there are sometimes multiple updates to the main branch per day and it takes the scientific tests about a week to run, many revisions in the main branch remain untested. In case of a test failure, the revision daemon performs a binary search bisecting the untested revisions to determine the exact revision that is responsible for the breakage.

[4.-N.] ***Testing daemons***. The testing daemons run on various platforms: Mac, Linux, and Windows. We currently have 18 of these daemons, some of which are meant for build tests (i.e., on Windows) and some of which are capable of running tests on our HPC cluster. Each daemon periodically checks the list of queued tests from the database server. If there is any test which that daemon is capable of running, it runs the test and then uploads the test results (logs, result files and test results encoded in JSON) to our SQL database.

This backend code is specific to our hardware, HPC use patterns and system administration environment, and maintained separately from the code that performs or tests science. This code does not include the frontend scientific testing framework (next paragraph) and is not needed to replicate any of the scientific results. The frontend implementation of the scientific testing framework including all of the scientific benchmarks are fully available under the RosettaCommons license.

### Setup of the scientific tests

We chose a simple setup as shown in FIG 1C. Each scientific test requires a small number of files, available in a template directory. All files in this directory are well documented with comments, and the lines that require editing for specific tests are highlighted. Each scientific test directory starts from a template containing the following files:

- *input files* – are either located in this directory or in a parallel git submodule if the input files exceed 5 MB. This policy prevents our main code repository from becoming overly inflated with thousands of input files for scientific benchmarking.
- *0.compile.py* – compiles the Rosetta and/or PyRosetta executable.
- *1.submit.py* – submits the benchmark jobs either to the local machine *or* to the HPC cluster. Note that this “or” provides hardware non-specificity; the user writes and debugs locally and can run at scale on the benchmark server.
- *2.analyze.py* – analyzes the output data, depending on the scientific objective. Analysis functions that are specific to this particular test live in this script, while broadly useful analysis functions are located in a file that is part of the overall Python test server framework and that contains functions to evaluate quality measures.
- *3.plot.py* – plots the output data via *matplotlib*^78^, or other plotting software as appropriate.
- … – other numbered scripts can be added as needed; they will run consecutively as numbered.
- *9.finalize.py* – gathers the output data and classifies the test as passed or failed, creates an HTML page by combining the documentation from the readme file, the plots of the output data and the pass/fail criterion. The HTML page is the main results page that developers, maintainers, and observers examine.
- *citation* – includes all relevant citations that describe the protocol, the benchmark set or the quality measures.
- *cutoffs* – contains the cutoffs used for distinguishing between a pass or a fail for this test.
- *observers* – email addresses of developers that either set up the test and / or maintain it. If a test fails on the test server, an email is sent to the observers to inform them of the test breakage.
- *readme.md* – is a questionnaire-style markdown file that contains all necessary documentation to understand the purpose and detailed methods of the test. Obtaining detailed documentation is essential for maintenance and longevity of the test. The goal is that anyone with basic Rosetta expertise and training can understand, reproduce, and maintain the test. The template readme file is provided in the supplement of this paper.

## Acknowledgements

ARO MURI W911NF-16-1-0372 to Watkins; American Heart Association 18POST34080422 to Kuenze; BSF 2015207 to Schueler-Furman, Ben-Aharon; Cancer Research Institute Irvington Postdoctoral Fellowship (CRI 3442) to Roy Burman; Candian Institutes of Health Research Postdoctoral Fellowship to Yachnin; Cyrus Biotechnology to Lewis; Simons Foundation to Bonneau, Koehler Leman, Mulligan; German Research Foundation KU 3510/1-1 to Kuenze; H2020 MSCA IF CC-LEGO 792305 to Ljubetic; HHMI to Baker; Hertz Foundation Fellowship to Alford; ISF 717/2017 to Schueler-Furman, Ben-Aharon; Lundbeck Foundation Fellowship R272-2017-4528 to Stein; Mistletoe Research Foundation Fellowship to Yachnin; NCN 2018/29/B/ST6/01989 to Gront, Krys; NIAID R01AI113867 to Schief, Adolf-Bryfogle; NIEHS P42ES004699 to Siegel; NIH 1R01GM123089 to Farrell, DiMaio; NIH 2R01GM098977 to Bailey-Kellogg; NIH F31-CA243353 to Smith; NIH F31-GM123616 to Jeliazkov; NIH GM067553 to Maguire; NIH NCI R21 CA219847 and NIH R01 GM121487 to Das, Watkins; NIH NHLBI 2R01HL128537 to Yarov-Yarovoy; NIH NIAID R21 AI156570 and NIH NIBIB R21 EB028342 to Bahl; NIH NIAID U01 AI150739, NIH NIDA R01 DA046138 and NIH NIGMS R01 GM080403 to Meiler; NIH NIGMS R01 GM073151 to Kuhlman, Gray, Leaver-Fay, Lyskov, Moretti, Meiler; NIH NIGMS R01 GM121487 and NIH NIGMS R35 GM122579 to Das; NIH NIGMS 1R01GM132110 and NIH NINDS 1R01NS103954 to Yarov-Yarovoy; NIH NINDS UG3NS114956 to Nguyen, Yarov-Yarovoy; NIH R01 DA046138 to Moretti; NIH F32 CA189246 to Labonte; NIH R01 GM 076324-11 to Siegel; NIH R01 GM080403 to Kuenze; NIH R01 GM129261 to Woods; NIH R01 GM078221 to Harmalkar, Roy Burman, Jeliazkov and Gray; NIH R01 GM127578 to Gray and Labonte; NIH R01 GM110089 to Loshbaugh, Kortemme, Barlow; NIH R35 GM131923 to Leaver-Fay, Teets, Kuhlman; NIH R01 GM132565 to Hansen, Khare; NSF 1507736 to Gray, Roy Burman; NSF 1627539 and NSF 1827246 to Siegel; NSF 1805510 to Siegel, Fell; NSF 2031785 to Bahl; NSF DBI-1564692 to Loshbaugh, Kortemme, Barlow and O’Connor; NSF GRFP Fellowship to Alford; NSF CBET1923691 to Hansen, Khare; Novo Nordisk Foundation NNF18OC0033950 to Tiemann, Stein, Lindorff-Larsen; RosettaCommons Licensing Fund RC8010 to Bahl; RosettaCommons to Hansen, Moretti, Lyskov, Khare, Gray; NIH NRSA T32AI007244 and NIH U19AI117905 to Schief, Adolf-Bryfogle.

The authors further thank Matt Mulqueen for expert administration of the multiple benchmark testing servers and cluster, RosettaCommons for hardware and staff support after the NIH ended their software infrastructure program, and companies that license Rosetta, providing support for critical software sustainability practices.

## Author Contributions

The benchmark testing server framework was implemented and is being maintained by SL. The scientific testing framework was created jointly by JKL, SL, and SML. Specific tests were implemented and validated by the test authors as outlined in Table 1. All tests went through independent scientific and technical review by SML, JKL, SL with help from VKM, RM, AMW and others for review of pull requests. Further, benchmarks were provided by JM, CB, KB, SOC, GK, and HW and independently reviewed by JKST, AS, KLL. JJG supervised the creation of the benchmark infrastructure and secured funding, together with BK. This project was jointly supervised by RB, JJG, DG, ALF, CB, CBK, DB, RD, FDM, SK, TK, JWL, JM, WS, OSF, JS, AS, VYY, and BK.

## Data availability

The frontend implementation of the scientific testing framework including all of the scientific benchmarks are fully available under the RosettaCommons license. Additionally, complete protocol captures for all benchmarks with input files, command lines, output files, analyses and result summaries are publicly available to view and download at https://graylab.jhu.edu/download/rosetta-scientific-tests/. These will be automatically expanded with new revisions about every 6 months. Older revisions remain on the server.

## References

1. Baker, M. & Penny, D. Is there a reproducibility crisis? Nature 533, 452–454 (2016).

2. Freedman, L. P., Cockburn, I. M. & Simcoe, T. S. The Economics of Reproducibility in Preclinical Research. PLOS Biol. 13, e1002165 (2015).

3. Peng, R. D. Reproducible research in computational science. Science (80-.). 334, 1226–1227 (2011).

4. Koehler Leman, J., Weitzner, B. D., Renfrew, P. D., Lewis, S. M., Moretti, R., Watkins, A. M., Mulligan, V. K., Lyskov, S., Adolf-Bryfogle, J., Labonte, J. W., Krys, J., Bystroff, C., Schief, W., Gront, D., Schueler-Furman, O., Baker, D., Bradley, P., Dunbrack, R., Kortemme, T., Leaver-Fay, A., Strauss, C. E. M., Meiler, J., Kuhlman, B., Gray, J. J. & Bonneau, R. Better together: Elements of successful scientific software development in a distributed collaborative community. PLOS Comput. Biol. 16, e1007507 (2020).

5. Adorf, C. S., Ramasubramani, V., Anderson, J. A. & Glotzer, S. C. How to professionally develop reusable scientific software-And when not to. Comput. Sci. Eng. 21, 66–79 (2019).

6. Baker, M. 1,500 scientists lift the lid on reproducibility: Nature News & Comment. Nature 533, 452 (2016).

7. Open Science Collaboration. Estimating the reproducibility of psychological science. Science (80-.). 349, aac4716–aac4716 (2015).

8. Stodden, V., Mcnutt, M., Bailey, D. H., Deelman, E., Gil, Y., Hanson, B., Heroux, M. A., Ioannidis, J. P. A. & Taufer, M. Enhancing reproducibility for computational methods. Science (80-.). 354, 1240–41 (2016).

9. Jeffrey Mervis. NSF to Ask Every Grant Applicant for Data Management Plan | Science | AAAS. Science (80-.). (2010). at <https://www.sciencemag.org/news/2010/05/nsf-ask-every-grant-applicant-data-management-plan>

10. Editorial. Everyone needs a data-management plan. Nature 555, 286–286 (2018).

11. Williams, M., Bagwell, J. & Nahm Zozus, M. Data management plans: the missing perspective. J. Biomed. Inform. 71, 130–142 (2017).

12. Wilkinson, M. D., Dumontier, M., Aalbersberg, Ij. J., Appleton, G., Axton, M., Baak, A., Blomberg, N., Boiten, J. W., da Silva Santos, L. B., Bourne, P. E., Bouwman, J., Brookes, A. J., Clark, T., Crosas, M., Dillo, I., Dumon, O., Edmunds, S., Evelo, C. T., Finkers, R., Gonzalez-Beltran, A., Gray, A. J. G., Groth, P., Goble, C., Grethe, J. S., Heringa, J., t Hoen, P. A. C., Hooft, R., Kuhn, T., Kok, R., Kok, J., Lusher, S. J., Martone, M. E., Mons, A., Packer, A. L., Persson, B., Rocca-Serra, P., Roos, M., van Schaik, R., Sansone, S. A., Schultes, E., Sengstag, T., Slater, T., Strawn, G., Swertz, M. A., Thompson, M., Van Der Lei, J., Van Mulligen, E., Velterop, J., Waagmeester, A., Wittenburg, P., Wolstencroft, K., Zhao, J. & Mons, B. The FAIR Guiding Principles for scientific data management and stewardship. Sci. Data 3, 1–9 (2016).

13. Afgan, E., Baker, D., van den Beek, M., Blankenberg, D., Bouvier, D., Čech, M., Chilton, J., Clements, D., Coraor, N., Eberhard, C., Grüning, B., Guerler, A., Hillman-Jackson, J., Von Kuster, G., Rasche, E., Soranzo, N., Turaga, N., Taylor, J., Nekrutenko, A. & Goecks, J. The Galaxy platform for accessible, reproducible and collaborative biomedical analyses: 2016 update. Nucleic Acids Res. 44, W3–W10 (2016).

14. Perkel, J. M. Challenge to scientists: does your ten-year-old code still run? Nature 584, 656–658 (2020).

15. ReScience C - Ten Years Reproducibility Challenge. at <https://rescience.github.io/ten-years/>

16. ReScience C. at <http://rescience.github.io/>

17. Van Bavel, J. J., Mende-Siedlecki, P., Brady, W. J. & Reinero, D. A. Contextual sensitivity in scientific reproducibility. Proc. Natl. Acad. Sci. 113, 6454–6459 (2016).

18. Peters, B., Brenner, S. E., Wang, E., Slonim, D. & Kann, M. G. Putting benchmarks in their rightful place: The heart of computational biology. PLOS Comput. Biol. 14, e1006494 (2018).

19. Ó Conchúir, S., Barlow, K. A., Pache, R. A., Ollikainen, N., Kundert, K., O’Meara, M. J., Smith, C. A. & Kortemme, T. A Web Resource for Standardized Benchmark Datasets, Metrics, and Rosetta Protocols for Macromolecular Modeling and Design. PLoS One 10, e0130433

20. Huizinga, D. & Kolawa, A. Automated Defect Prevention: Best Practices in Software Management |Wiley. 2007). at <https://www.wiley.com/en-us/Automated+Defect+Prevention%3A+Best+Practices+in+Software+Management-p-9780470042120>

21. Moult, J., Fidelis, K., Kryshtafovych, A., Schwede, T. & Tramontano, A. Critical assessment of methods of protein structure prediction (CASP)-Round XII. Proteins Struct. Funct. Bioinforma. 86, 7–15 (2018).

22. Wodak, S. J. & Janin, J. Modeling protein assemblies: Critical Assessment of Predicted Interactions (CAPRI) 15 years hence. Proteins Struct. Funct. Bioinforma. 85, 357–358 (2017).

23. Friedberg, I. & Radivojac, P. in Methods Mol. Biol. 1446, 133–146 (2017).

24. Daneshjou, R., Wang, Y., Bromberg, Y., Bovo, S., Martelli, P. L., Babbi, G., Lena, P. Di, Casadio, R., Edwards, M., Gifford, D., Jones, D. T., Sundaram, L., Bhat, R. R., Li, X., Pal, L. R., Kundu, K., Yin, Y., Moult, J., Jiang, Y., Pejaver, V., Pagel, K. A., Li, B., Mooney, S. D., Radivojac, P., Shah, S., Carraro, M., Gasparini, A., Leonardi, E., Giollo, M., Ferrari, C., Tosatto, S. C. E., Bachar, E., Azaria, J. R., Ofran, Y., Unger, R., Niroula, A., Vihinen, M., Chang, B., Wang, M. H., Franke, A., Petersen, B.-S., Pirooznia, M., Zandi, P., McCombie, R., Potash, J. B., Altman, R. B., Klein, T. E., Hoskins, R. A., Repo, S., Brenner, S. E. & Morgan, A. A. Working toward precision medicine: Predicting phenotypes from exomes in the Critical Assessment of Genome Interpretation (CAGI) challenges. Hum. Mutat. 38, 1182–1192 (2017).

25. Miao, Z., Adamiak, R. W., Antczak, M., Boniecki, M. J., Bujnicki, J., Chen, S. J., Cheng, C. Y., Cheng, Y., Chou, F. C., Das, R., Dokholyan, N. V., Ding, F., Geniesse, C., Jiang, Y., Joshi, A., Krokhotin, A., Magnus, M., Mailhot, O., Major, F., Mann, T. H., Piątkowski, P., Pluta, R., Popenda, M., Sarzynska, J., Sun, L., Szachniuk, M., Tian, S., Wang, J., Wang, J., Watkins, A. M., Wiedemann, J., Xiao, Y., Xu, X., Yesselman, J. D., Zhang, D., Zhang, Y., Zhang, Z., Zhao, C., Zhao, P., Zhou, Y., Zok, T., Żyła, A., Ren, A., Batey, R. T., Golden, B. L., Huang, L., Lilley, D. M., Liu, Y., Patel, D. J. & Westhof, E. RNA-Puzzles round IV: 3D Structure predictions of four ribozymes and two aptamers. RNA 26, (2020).

26. Haas, J., Barbato, A., Behringer, D., Studer, G., Roth, S., Bertoni, M., Mostaguir, K., Gumienny, R. & Schwede, T. Continuous Automated Model EvaluatiOn (CAMEO) complementing the critical assessment of structure prediction in CASP12. Proteins Struct. Funct. Bioinforma. 86, 387–398 (2018).

27. Leaver-Fay, A., Tyka, M., Lewis, S. M., Lange, O. F., Thompson, J. M., Jacak, R., Kaufman, K., Renfrew, P. D., Smith, C. A., Sheffler, W., Davis, I. W., Cooper, S., Treuille, A., Mandell, D. J., Richter, F., Ban, Y.-E. A., Fleishman, S. J., Corn, J. E., Kim, D. E., Berrondo, M., Mentzer, S., Popovic, Z., Havranek, J. J., Karanicolas, J., Das, R., Meiler, J., Kortemme, T., Gray, J. J., Kuhlman, B., Baker, D. & Bradley, P. ROSETTA3: An Object-Oriented Software Suite for the Simulation and Design of Macromolecules. Methods Enzymol. 487, 545–74 (2011).

28. Koehler Leman, J., Weitzner, B. D., Lewis, S. M., Adolf-Bryfogle, J., Alam, N., Alford, R. F., Aprahamian, M., Baker, D., Barlow, K. A., Barth, P., Basanta, B., Bender, B. J., Blacklock, K., Bonet, J., Boyken, S. E., Bradley, P., Bystroff, C., Conway, P., Cooper, S., Correia, B. E., Coventry, B., Das, R., De Jong, R. M., DiMaio, F., Dsilva, L., Dunbrack, R., Ford, A. S., Frenz, B., Fu, D. Y., Geniesse, C., Goldschmidt, L., Gowthaman, R., Gray, J. J., Gront, D., Guffy, S., Horowitz, S., Huang, P. S., Huber, T., Jacobs, T. M., Jeliazkov, J. R., Johnson, D. K., Kappel, K., Karanicolas, J., Khakzad, H., Khar, K. R., Khare, S. D., Khatib, F., Khramushin, A., King, I. C., Kleffner, R., Koepnick, B., Kortemme, T., Kuenze, G., Kuhlman, B., Kuroda, D., Labonte, J. W., Lai, J. K., Lapidoth, G., Leaver-Fay, A., Lindert, S., Linsky, T., London, N., Lubin, J. H., Lyskov, S., Maguire, J., Malmström, L., Marcos, E., Marcu, O., Marze, N. A., Meiler, J., Moretti, R., Mulligan, V. K., Nerli, S., Norn, C., Ó’Conchúir, S., Ollikainen, N., Ovchinnikov, S., Pacella, M. S., Pan, X., Park, H., Pavlovicz, R. E., Pethe, M., Pierce, B. G., Pilla, K. B., Raveh, B., Renfrew, P. D., Burman, S. S. R., Rubenstein, A., Sauer, M. F., Scheck, A., Schief, W., Schueler-Furman, O., Sedan, Y., Sevy, A. M., Sgourakis, N. G., Shi, L., Siegel, J. B., Silva, D. A., Smith, S., Song, Y., Stein, A., Szegedy, M., Teets, F. D., Thyme, S. B., Wang, R. Y. R., Watkins, A., Zimmerman, L. & Bonneau, R. Macromolecular modeling and design in Rosetta: recent methods and frameworks. Nat. Methods 17, 665–680 (2020).

29. RosettaCommons. at <https://www.rosettacommons.org/>

30. Kaufmann, K. W. & Meiler, J. Using RosettaLigand for Small Molecule Docking into Comparative Models. PLoS One 7, e50769 (2012).

31. Conway, P., Tyka, M. D., DiMaio, F., Konerding, D. E. & Baker, D. Relaxation of backbone bond geometry improves protein energy landscape modeling. Protein Sci. 23, 47–55 (2014).

32. Leaver-Fay, A., O’Meara, M. J., Tyka, M., Jacak, R., Song, Y., Kellogg, E. H., Thompson, J., Davis, I. W., Pache, R. A., Lyskov, S., Gray, J. J., Kortemme, T., Richardson, J. S., Havranek, J. J., Snoeyink, J., Baker, D. & Kuhlman, B. Scientific benchmarks for guiding macromolecular energy function improvement. Methods Enzymol. 523, 109–43 (2013).

33. O’Meara, M. J., Leaver-Fay, A., Tyka, M. D., Stein, A., Houlihan, K., DiMaio, F., Bradley, P., Kortemme, T., Baker, D., Snoeyink, J. & Kuhlman, B. Combined Covalent-Electrostatic Model of Hydrogen Bonding Improves Structure Prediction with Rosetta. J. Chem. Theory Comput. 11, 609–622 (2015).

34. Park, H., Bradley, P., Greisen, P., Liu, Y., Mulligan, V. K., Kim, D. E., Baker, D. & DiMaio, F. Simultaneous Optimization of Biomolecular Energy Functions on Features from Small Molecules and Macromolecules. J. Chem. Theory Comput. 12, 6201–6212 (2016).

35. Alford, R. F., Leaver-Fay, A., Jeliazkov, J. R., O’Meara, M. J., Dimaio, F. P., Park, H., Shapovalov, M. V., Renfrew, P. D., Mulligan, V. K., Kappel, K., Labonte, J. W., Pacella, M. S., Bonneau, R., Bradley, P., Dunbrack, R. L., Das, R., Baker, D., Kuhlman, B., Kortemme, T. & Gray, J. J. The Rosetta all-atom energy function for macromolecular modeling and design. J. Chem. Theory Comput. 13, 1–35 (2017).

36. Alford, R. F. & Gray, J. J. Diverse scientific benchmarks for implicit membrane energy functions. bioRxiv [Preprint] 2020.06.23.168021 (2020). doi:10.1101/2020.06.23.168021

37. Renfrew, P. D., Campbell, G., Strauss, C. E. M. & Bonneau, R. The 2010 Rosetta Developers Meeting: Macromolecular Prediction and Design Meets Reproducible Publishing. PLoS One 6, e22431 (2011).

38. Bender, B. J., Cisneros, A., Duran, A. M., Finn, J. A., Fu, D., Lokits, A. D., Mueller, B. K., Sangha, A. K., Sauer, M. F., Sevy, A. M., Sliwoski, G., Sheehan, J. H., Dimaio, F., Meiler, J. & Moretti, R. Protocols for Molecular Modeling with Rosetta3 and RosettaScripts. Biochemistry acs.biochem.6b00444 (2016). doi:10.1021/acs.biochem.6b00444

39. Fleishman, S. J., Leaver-Fay, A., Corn, J. E., Strauch, E.-M. M., Khare, S. D., Koga, N., Ashworth, J., Murphy, P., Richter, F., Lemmon, G., Meiler, J. & Baker, D. RosettaScripts: A scripting language interface to the Rosetta Macromolecular modeling suite. PLoS One 6, 1–10 (2011).

40. Chaudhury, S., Lyskov, S. & Gray, J. J. PyRosetta: a script-based interface for implementing molecular modeling algorithms using Rosetta. Bioinformatics 26, 689–691 (2010).

41. Gray, J. J., Chaudhury, S., Lyskov, S., and Labonte, J. W. The PyRosetta Interactive Platform for Protein Structure Prediction and Design: A Set of Educational Modules. (2014). at <http://www.amazon.com/PyRosetta-Interactive-Platform-Structure-Prediction/dp/1500968277>

42. RosettaCommons. Rosetta documentation - Scientific Benchmarks. at <http://new.rosettacommons.org/docs/latest/development_documentation/test/Scientific-Benchmarks>

43. Weitzner, B. D., Jeliazkov, J. R., Lyskov, S., Marze, N., Kuroda, D., Frick, R., Adolf-Bryfogle, J., Biswas, N., Dunbrack, R. L. & Gray, J. J. Modeling and docking of antibody structures with Rosetta. Nat. Protoc. 12, 401–416 (2017).

44. Weitzner, B. D. & Gray, J. J. Accurate Structure Prediction of CDR H3 Loops Enabled by a Novel Structure-Based C-Terminal Constraint. J. Immunol. 198, 505–515 (2017).

45. Sircar, A. & Gray, J. J. SnugDock: paratope structural optimization during antibody-antigen docking compensates for errors in antibody homology models. PLoS Comput. Biol. 6, e1000644 (2010).

46. Labonte, J. W., Adolf-Bryfogle, J., Schief, W. R. & Gray, J. J. Residue-centric modeling and design of saccharide and glycoconjugate structures. J. Comput. Chem. 38, 276–287 (2017).

47. Adolf-Bryfogle, J., Labonte, J. W., Kraft, J., Shapovalov, M. V, Raemisch, S., Luettke, T., DiMaio, F., Bahl, C. D., Palleson, J., King, N. P., Gray, J. J., Kulp, D. W. & Schief, W. R. Growing Glycans in Rosetta: Accurate de-novo glycan modeling, density fitting, and rational sequon design. Prep. (2021).

48. Song, Y., Dimaio, F., Wang, R. Y.-R. R., Kim, D. E., Miles, C., Brunette, T., Thompson, J. & Baker, D. High-resolution comparative modeling with RosettaCM. Structure 21, 1735–1742 (2013).

49. Kortemme, T., Kim, D. E. & Baker, D. Computational alanine scanning of protein-protein interfaces. Sci. STKE 2004, pl2 (2004).

50. Guffy, S. L., Teets, F. D., Langlois, M. I. & Kuhlman, B. Protocols for Requirement-Driven Protein Design in the Rosetta Modeling Program. J. Chem. Inf. Model. 58, 895–901 (2018).

51. Nivón, L. G., Bjelic, S., King, C. & Baker, D. Automating human intuition for protein design. Proteins 82, 858–66 (2014).

52. Maguire, J. B., Haddox, H. K., Strickland, D., Halabiya, S. F., Coventry, B., Griffin, J. R., Pulavarti, S. V. S. R. K., Cummins, M., Thieker, D. F., Klavins, E., Szyperski, T., DiMaio, F., Baker, D. & Kuhlman, B. Perturbing the energy landscape for improved packing during computational protein design. Proteins Struct. Funct. Bioinforma. 89, 436–449 (2021).

53. Yachnin, B. J., Mulligan, V. K., Khare, S. D. & Bailey-Kellogg, C. MHCEpitopeEnergy, a flexible Rosetta-based biotherapeutic deimmunization platform. J. Chem. Inf. Model. in revision, (2021).

54. Gray, J. J., Moughon, S., Wang, C., Schueler-Furman, O., Kuhlman, B., Rohl, C. A. & Baker, D. Protein–Protein Docking with Simultaneous Optimization of Rigid-body Displacement and Side-chain Conformations. J. Mol. Biol. 331, 281–99 (2003).

55. Marze, N. A., Roy Burman, S. S., Sheffler, W. & Gray, J. J. Efficient flexible backbone protein–protein docking for challenging targets. Bioinformatics 34, 3461–3469 (2018).

56. Alam, N. & Schueler-Furman, O. in Methods Mol. Biol. 1561, 139–169 (Humana Press Inc., 2017).

57. Gront, D., Kulp, D. W., Vernon, R. M., Strauss, C. E. M. & Baker, D. Generalized Fragment Picking in Rosetta : Design, Protocols and Applications. 6, (2011).

58. Loshbaugh, A. L. & Kortemme, T. Comparison of Rosetta flexible-backbone computational protein design methods on binding interactions. Proteins Struct. Funct. Bioinforma. 88, 206–226 (2020).

59. Smith, S. T. & Meiler, J. Assessing multiple score functions in Rosetta for drug discovery. PLoS One 15, e0240450 (2020).

60. Canutescu, A. A. & Dunbrack, R. L. Cyclic coordinate descent: A robotics algorithm for protein loop closure. Protein Sci. 12, 963–72 (2003).

61. Mandell, D. J., Coutsias, E. A. & Kortemme, T. Sub-angstrom accuracy in protein loop reconstruction by robotics-inspired conformational sampling. Nat. Methods 6, 551–2 (2009).

62. Fernandez, A. J., Daniel, E. J. P., Mahajan, S. P., Gray, J. J., Gerken, T. A., Tabak, L. A. & Samara, N. L. The structure of the colorectal cancer-associated enzyme GalNAc-T12 reveals how nonconserved residues dictate its function. Proc. Natl. Acad. Sci. U. S. A. 116, 20404–20410 (2019).

63. Stein, A. & Kortemme, T. Improvements to robotics-inspired conformational sampling in rosetta. PLoS One 8, e63090 (2013).

64. Alford, R. F., Fleming, P. J., Fleming, K. G. & Gray, J. J. Protein Structure Prediction and Design in a Biologically Realistic Implicit Membrane. Biophys. J. 118, 2042–2055 (2020).

65. Alford, R. F., Koehler Leman, J., Weitzner, B. D., Duran, A. M., Tilley, D. C., Elazar, A. & Gray, J. J. An Integrated Framework Advancing Membrane Protein Modeling and Design. PLoS Comput. Biol. 11, e1004398 (2015).

66. Koehler Leman, J. & Bonneau, R. A Novel Domain Assembly Routine for Creating Full-Length Models of Membrane Proteins from Known Domain Structures. Biochemistry 57, 1939–1944 (2018).

67. Koehler Leman, J., Lyskov, S. & Bonneau, R. Computing structure-based lipid accessibility of membrane proteins with mp_lipid_acc in RosettaMP. BMC Bioinformatics 18, 115 (2017).

68. Bhardwaj, G., Mulligan, V. K., Bahl, C. D., Gilmore, J. M., Harvey, P. J., Cheneval, O., Buchko, G. W., Pulavarti, S. V. S. R. K., Kaas, Q., Eletsky, A., Huang, P.-S., Johnsen, W. A., Greisen, P. J., Rocklin, G. J., Song, Y., Linsky, T. W., Watkins, A., Rettie, S. A., Xu, X., Carter, L. P., Bonneau, R., Olson, J. M., Coutsias, E., Correnti, C. E., Szyperski, T., Craik, D. J. & Baker, D. Accurate de novo design of hyperstable constrained peptides. Nature 538, 329–335 (2016).

69. Mulligan, V. K., Workman, S., Sun, T., Rettie, S., Li, X., Worrall, L. J., Craven, T. W., King, D. T., Hosseinzadeh, P., Watkins, A. M., Douglas Renfrew, P., Guffy, S., Labonte, J. W., Moretti, R., Bonneau, R., Strynadka, N. C. J. & Baker, D. Computationally designed peptide macrocycle inhibitors of New Delhi metallo-β-lactamase 1. Proc. Natl. Acad. Sci. U. S. A. 118, (2021).

70. Tyka, M. D., Keedy, D. A., André, I., DiMaio, F., Song, Y., Richardson, D. C., Richardson, J. S. & Baker, D. Alternate States of Proteins Revealed by Detailed Energy Landscape Mapping. J. Mol. Biol. 405, 607–618 (2011).

71. Watkins, A. M., Rangan, R. & Das, R. FARFAR2: Improved De Novo Rosetta Prediction of Complex Global RNA Folds. Structure 28, 963–976.e6 (2020).

72. Watkins, A. M., Geniesse, C., Kladwang, W., Zakrevsky, P., Jaeger, L. & Das, R. Blind prediction of noncanonical RNA structure at atomic accuracy. Sci. Adv. 4, eaar5316 (2018).

73. Hosseinzadeh, P., Bhardwaj, G., Mulligan, V. K., Shortridge, M. D., Craven, T. W., Pardo-Avila, F., Rettie, S. A., Kim, D. E., Silva, D.-A., Ibrahim, Y. M., Webb, I. K., Cort, J. R., Adkins, J. N., Varani, G. & Baker, D. Comprehensive computational design of ordered peptide macrocycles. Science (80-.). 358, 1461–1466 (2017).

74. Travis CI - continuous integration. https://travis-ci.org/

75. Jenkins. https://jenkins.io/

76. Lyskov, S., Chou, F.-C., Conchúir, S. Ó., Der, B. S., Drew, K., Kuroda, D., Xu, J., Weitzner, B. D., Renfrew, P. D., Sripakdeevong, P., Borgo, B., Havranek, J. J., Kuhlman, B., Kortemme, T., Bonneau, R., Gray, J. J. & Das, R. Serverification of molecular modeling applications: the Rosetta Online Server that Includes Everyone (ROSIE). PLoS One 8, e63906 (2013).

77. GitHub. https://github.com/

78. Matplotlib: Python plotting — Matplotlib 3.4.1 documentation. at <https://matplotlib.org/>

