## Supplement for "Ensuring scientific reproducibility in bio-macromolecular modeling via extensive, automated benchmarks"

*Documentation template in markdown format*

### AUTHOR AND DATE
##### Who set up the benchmark? Please add name, email, PI, month and year

### PURPOSE OF THE TEST
##### What does the benchmark test and why?

### BENCHMARK DATASET
##### How many proteins are in the set?
##### What dataset are you using? Is it published? If yes, please add a citation.
##### What are the input files? How were they created?

### PROTOCOL
##### State and briefly describe the protocol.
##### Is there a publication that describes the protocol?
##### How many CPU hours does this benchmark take approximately?

### PERFORMANCE METRICS
##### What are the performance metrics used and why were they chosen?
##### How do you define a pass/fail for this test?
##### How were any cutoffs defined?

### KEY RESULTS
##### What is the baseline to compare things to - experimental data or a previous Rosetta protocol?
##### Describe outliers in the dataset.

### DEFINITIONS AND COMMENTS
##### State anything you think is important for someone else to replicate your results.

### LIMITATIONS
##### What are the limitations of the benchmark? Consider dataset, quality measures, protocol etc.
##### How could the benchmark be improved?
##### What goals should be hit to make this a "good" benchmark?

*Notes on our test server framework*

We use our test server architecture to run unit, integration, build, and performance tests. The integration between hardware and software in the test server allows anyone in our community to schedule any tests for any Rosetta development branches (their own, or other community members') that have been pushed to the version control server GitHub[1]. The test server web interface (FIG. 2A; available at [https://benchmark.graylab.jhu.edu/](https://benchmark.graylab.jhu.edu/)) is integrated with GitHub and its pull-request feature allows scheduling of tests via branch name, pull-request identifier, or GitHub commit *sha1* key. Dedicated maintainers ('observers') are notified of test breakage via email and community members can view all test results on the test server web interface after GitHub login. The HPC cluster that runs these computations is partly funded through government grants and funds from the RosettaCommons[2].

*Steps to add a scientific test to the Rosetta test server framework for continuous, automated testing*

Once the dataset, interface (command line, RosettaScripts, or PyRosetta), specific command line, and quality measures have been chosen, the author can use the template and follow the steps outlined on the documentation page[3] to contribute the test.

The test should be created in a feature branch, pushed to GitHub and a pull-request should be created for review. Once the test runs without errors locally, it can be run on the test server. Having individually numbered scripts from compilation to finalizing the results page facilitates debugging. Once the test runs without errors locally and on the test server and reviewers approved, the feature branch is merged into Rosetta's main development branch. The test will then run 'continuously' from the 'oldest' (i.e., the test that was run longest ago) to the newest, based on node availability, and the results are stored in the database.

*Documentation: Defining appropriate cutoffs for pass/fail is crucial for longevity*

Scientific tests check the scientific performance of an entire protocol or workflow. Since Rosetta samples via a Monte-Carlo procedure, the randomness from the sampling will affect the results, making it more difficult to generalize a pass/fail criterion. Other difficulties are the varying scientific objectives of different tests and the question of how to set the cutoff for the test to be most useful and maintainable. For instance, if the cutoffs are such that the test almost never passes, the test will be ignored as fundamentally miswritten and becomes essentially meaningless. The goal is to find an optimal cutoff that probes scientific validity while failing only when the latter is compromised. We determine the cutoffs by defining specific quality measures, running each protocol several times, and adjusting the cutoffs such that the tests generally pass, but fail when the output shifts too far from its original position. We find that tests that define a cutoff by adding a small number of standard deviations to the mean of the results are generally useful.

*Benchmarks sets are diverse and provide a more realistic performance*

One of the problems with publishing newly developed or improved methods is the pressure to promote the method as best as possible to warrant publication and use by the scientific community. This is often at odds with presenting the method in a realistic light. The methods' performance is often artificially inflated through cherry picking targets and removing outliers, or its improvement is marginal and not statistically relevant. Overly positive presentations pertain to the collection of targets in the benchmark dataset, the choice of quality measures and the appropriate use of statistics. To produce a realistic performance of the method, we ask our authors to collect benchmark sets that prevent biases in the presentation of results and to choose examples that are diverse in prediction difficulty and other independent dimensions or features. Explicitly stating shortcomings and areas of improvement for the benchmark set, method, or quality measures directs new developers towards areas of improvements for future method development tasks.

*Score function implementations suffer from heterogeneity*

One possible area of improvement that the scientific benchmarks revealed is the different use and implementations of score functions across applications. The heterogeneity in score function implementation makes it impossible to easily test different score functions for all applications and hinders progress in development. Some applications use and implement hardcoded score functions, some use the one provided in the command line, some use the ones provided in the RosettaScripts interface. Some applications implement different score functions for low- and high-resolution modes, sometimes employing different ones for sampling and scoring. Some applications use special score functions, specifically designed for a particular application and improved over the years, sometimes for different stages of the protocol or with and without specific constraints or features turned on or off.

### References


1. GitHub. *https://github.com/*
2. Koehler Leman, J., Weitzner, B. D., Renfrew, P. D., Lewis, S. M., Moretti, R., Watkins, A. M., Mulligan, V. K., Lyskov, S., Adolf-Bryfogle, J., Labonte, J. W., Krys, J., Bystroff, C., Schief, W., Gront, D., Schueler-Furman, O., Baker, D., Bradley, P., Dunbrack, R., Kortemme, T., Leaver-Fay, A., Strauss, C. E. M., Meiler, J., Kuhlman, B., Gray, J. J. & Bonneau, R. Better together: Elements of successful scientific software development in a distributed collaborative community. *PLOS Comput. Biol.* **16,** e1007507 (2020).
3. RosettaCommons. Rosetta documentation - Scientific Benchmarks. at <http://new.rosettacommons.org/docs/latest/development_documentation/test/Scientific-Benchmarks>
